# Nucleotide Analogues as Inhibitors of SARS-CoV Polymerase

**DOI:** 10.1101/2020.03.12.989186

**Authors:** Jingyue Ju, Xiaoxu Li, Shiv Kumar, Steffen Jockusch, Minchen Chien, Chuanjuan Tao, Irina Morozova, Sergey Kalachikov, Robert N. Kirchdoerfer, James J. Russo

## Abstract

SARS-CoV-2, a member of the coronavirus family, has caused a global public health emergency.^1^ Based on our analysis of hepatitis C virus and coronavirus replication, and the molecular structures and activities of viral inhibitors, we previously reasoned that the FDA-approved heptatitis C drug EPCLUSA (Sofosbuvir/Velpatasvir) should inhibit coronaviruses, including SARS-CoV-2.^2^ Here, using model polymerase extension experiments, we demonstrate that the activated triphosphate form of Sofosbuvir is incorporated by low-fidelity polymerases and SARS-CoV RNA-dependent RNA polymerase (RdRp), and blocks further incorporation by these polymerases; the activated triphosphate form of Sofosbuvir is not incorporated by a host-like high-fidelity DNA polymerase. Using the same molecular insight, we selected two other anti-viral agents, Alovudine and AZT (an FDA approved HIV/AIDS drug) for evaluation as inhibitors of SARS-CoV RdRp. We demonstrate the ability of two HIV reverse transcriptase inhibitors, 3’-fluoro-3’-deoxythymidine triphosphate and 3’-azido-3’-deoxythymidine triphosphate (the active triphosphate forms of Alovudine and AZT), to be incorporated by SARS-CoV RdRp where they also terminate further polymerase extension. Given the 98% amino acid similarity of the SARS-CoV and SARS-CoV-2 RdRps, we expect these nucleotide analogues would also inhibit the SARS-CoV-2 polymerase. These results offer guidance to further modify these nucleotide analogues to generate more potent broad-spectrum anti-coronavirus agents.

---

The recent appearance of a new coronaviral infection, COVID-19, in Wuhan, China, and its worldwide spread, has made international headlines. Already more than 3,000 deaths have been ascribed to this virus, and COVID-19 has reached near pandemic status. The virus has been isolated from the lower respiratory tracts of patients with pneumonia, sequenced and visualized by electron microscopy.^1^ The virus, designated SARS-CoV-2, is a new member of the subgenus *Sarbecovirus*, in the Orthocoronavirinae subfamily, but is distinct from MERS-CoV and SARS-CoV.^1^ The coronaviruses are single strand RNA viruses, sharing properties with other single-stranded RNA viruses such as hepatitis C virus (HCV), West Nile virus, Marburg virus, HIV virus, Ebola virus, dengue virus, and rhinoviruses. In particular, coronaviruses and HCV are both positive-sense single-strand RNA viruses,^3,4^ and thus have a similar replication mechanism requiring a RNA-dependent RNA polymerase (RdRp).

The coronavirus life cycle has been described.^3^ Briefly, the virus enters the cell by endocytosis, is uncoated, and ORF1a and ORF1b of the positive strand RNA is translated to produce nonstructural protein precursors, including a cysteine protease and a serine protease; these further cleave the precursors to form mature, functional helicase and RNA-dependent RNA polymerase. A replication-transcription complex is then formed, which is responsible for making more copies of the RNA genome via a negative-sense RNA intermediate, as well as the structural and other proteins encoded by the viral genome. The viral RNA is packaged into viral coats in the endoplasmic reticulum-Golgi intermediate complex, after which exocytosis results in release of viral particles for subsequent infectious cycles. Potential inhibitors have been designed to target nearly every stage of this process.^3^ However, despite decades of research, no effective drug is currently approved to treat serious coronavirus infections such as SARS, MERS, and COVID-19.

One of the most important druggable targets for coronaviruses is the RNA-dependent RNA polymerase (RdRp). This polymerase is highly conserved at the protein level among different positive sense RNA viruses, to which coronaviruses and HCV belong, and shares common structural features in these viruses.^5^ Like RdRps in other viruses, the coronavirus enzyme is highly error-prone,^6^ which might increase its ability to accept modified nucleotide analogues. Nucleotide and nucleoside analogues that inhibit polymerases are an important group of anti-viral agents.^7–10^

Based on our analysis of hepatitis C virus and coronavirus replication, and the molecular structures and activities of viral inhibitors, we previously reasoned that the FDA-approved heptatitis C drug EPCLUSA (Sofosbuvir/Velpatasvir) should inhibit coronaviruses, including SARS-CoV-2.^2^ Sofosbuvir is a pyrimidine nucleotide analogue prodrug with a hydrophobic masked phosphate group enabling it to enter infected eukaryotic cells, and then converted into its active triphosphate form by cellular enzymes (Fig. 1). In this activated form, it inhibits the HCV RNA-dependent RNA polymerase NS5B.^11,12^ The activated drug (2’-F,Me-UTP) binds in the active site of the RdRp, where it is incorporated into RNA, and due to fluoro and methyl modifications at the 2’ position, inhibits further RNA chain extension, thereby halting RNA replication and stopping viral growth. It acts as an RNA polymerase inhibitor by competing with natural ribonucleotides. Velpatasvir inhibits NS5A, a key protein required for HCV replication. NS5A enhances the function of RNA polymerase NS5B during viral RNA synthesis.^13,14^

**Fig. 1.**
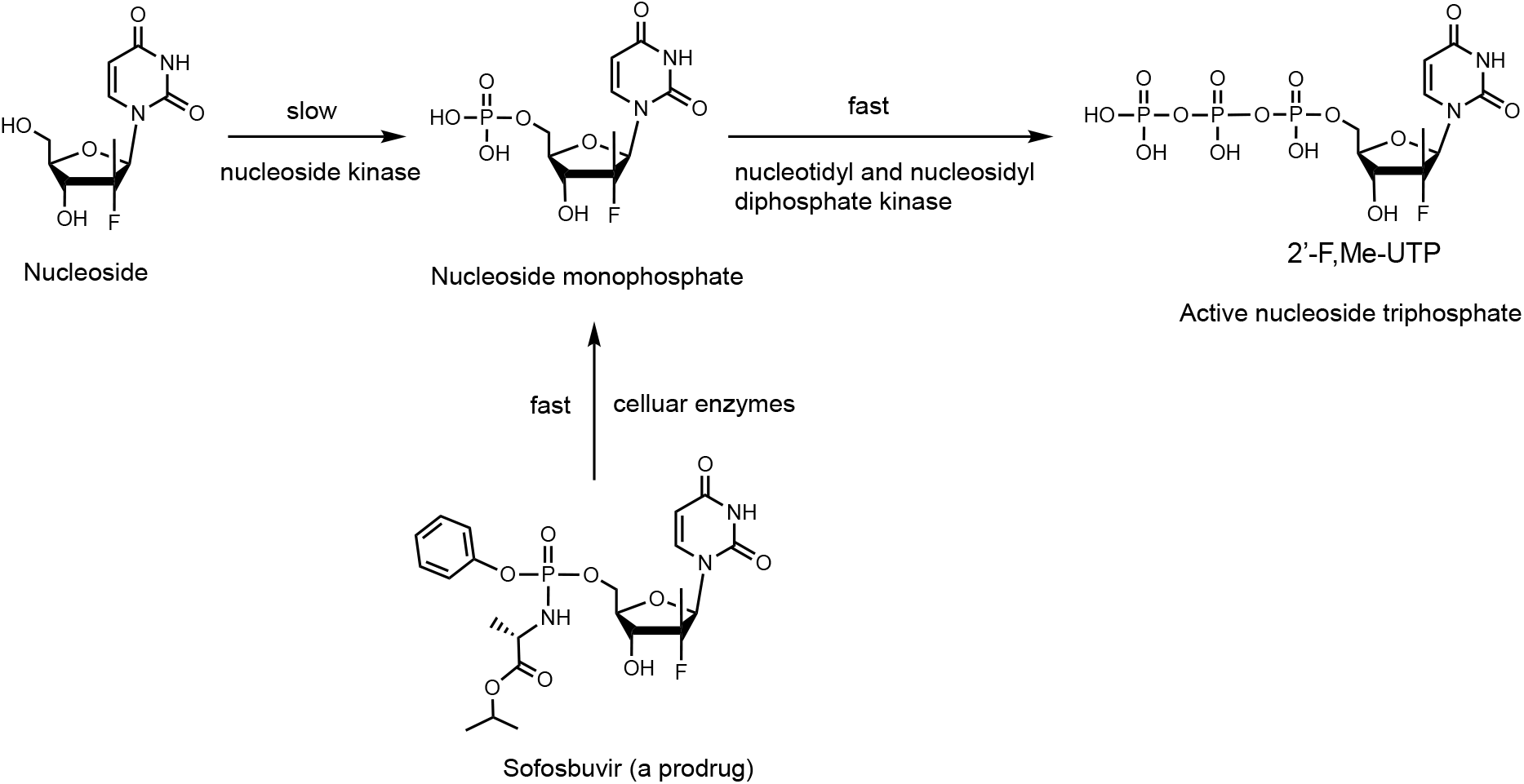
Conversion of Sofosbuvir to active triphosphate (2’-F,Me-UTP) *in vivo* to inhibit viral polymerases. *Adapted from* ^11^.

There are many other RNA polymerase inhibitors that have been evaluated as antiviral drugs. A related purine nucleotide prodrug, Remdesivir (Fig. 2b), was developed by Gilead to treat Ebola virus infections, though not successfully, and is currently in clinical trials for treating COVID-19.^15,16^ In contrast to Sofosbuvir (Fig. 2a), both the 2’- and 3’-OH groups in Remdesivir (Fig. 2b) are unmodified, but a cyano group at the 1’ position serves to inhibit the RdRp in the active triphosphate form. In addition to the use of hydrophobic groups to mask the phosphate in the Protide-based prodrug strategy,^17^ as with Sofosbuvir and Remdesivir, there are other classes of nucleoside prodrugs including those based on ester derivatives of the ribose hydroxyl groups to enhance cellular delivery.^18,19^

**Fig. 2.**
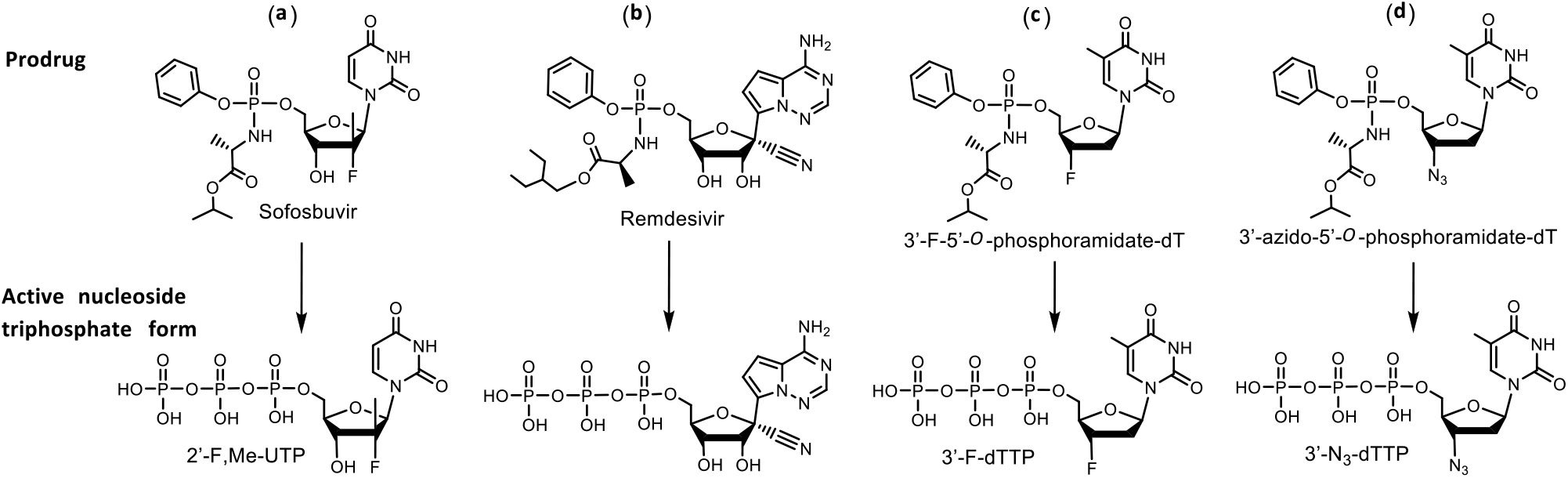
Comparison of structures of prodrug viral inhibitors. Top: Prodrug (phosphoramidate) form; Bottom: Active triphosphorylated form.

The replication cycle of HCV^4^ is very similar to that of the coronaviruses.^3^ Analyzing the structure of the active triphosphate form of Sofosbuvir (Fig. 2a) compared to that of Remdesivir (Fig. 2b), both of which have already been shown to inhibit the replication of specific RNA viruses (Sofosbuvir for HCV, Remdesivir for SARS-CoV-2), we noted in particular that the 2’-modifications in Sofosbuvir (a fluoro and a methyl group) are substantially smaller than the 1’-cyano group and the 2’-OH group in Remdesivir. The bulky cyano group in close proximity to the 2’-OH may lead to steric hindrance that will impact the incorporation efficiency of the activated form of Remdesivir. Interestingly, it was recently reported that, using the MERS-CoV polymerase, the triphosphate of Remdesivir was preferentially incorporated relative to ATP in solution assays.^20^ Nevertheless, it has been shown that the active triphosphate form of Remdesivir does not cause complete polymerase reaction termination and actually delays polymerase termination in Ebola virus and respiratory syncytial virus, likely due to its 1’-cyano group and the free 2’-OH and 3’-OH groups.^20,21^ Compared to the active form of Sofosbuvir (2’-fluoro-2’-methyl-UTP), two other nucleotide inhibitors with related structures were reviewed: 2’-fluoro-UTP is incorporated by polymerase, but RNA synthesis may continue past the incorporated nucleotide analogue;^22^ 2’-C-methyl-UTP has been shown to terminate the reaction catalyzed by HCV RdRp,^22^ but proofreading mechanisms can revert this inhibition in mitochondrial DNA-dependent RNA polymerases.^23^ Additionally, HCV develops resistance to 2’-C-methyl-UTP due to mutations of the RdRp.^24^ A computational study published in 2017^25^ considered the ability of various anti-HCV drugs to dock in the active site of SARS and MERS coronavirus RdRps as potential inhibitors. Recently, Elfiky used a computational approach to predict that Sofosbuvir, IDX-184, Ribavirin, and Remidisvir might be potent drugs against COVID-19.^26^

Thus, based on our analysis of the biological pathways of hepatitis C and coronaviruses, the molecular structures and activities of viral inhibitors, model polymerase and SARS-CoV RdRp extension experiments described below, and the efficacy of Sofosbuvir in inhibiting the HCV RdRp, we expect that Sofosbuvir or its modified forms should also inhibit the SARS-CoV-2 polymerase.^2^

The active triphosphate form of Sofosbuvir (2’-F,Me-UTP) was shown to be incorporated by HCV RdRp and prevent any further incorporation by this polymerase.^22,27^ Other viral polymerases have also been shown to incorporate active forms of various anti-viral prodrugs to inhibit replication.^28^ Since, at the time of the preparation of this manuscript, we did not have access to the RdRp from SARS-CoV-2, we first selected two groups of polymerases to test the termination efficiency of the active form of Sofosbuvir, one group with high fidelity behavior with regard to incorporation of modified nucleotide analogues, which one would expect for host cell polymerases, the other group with low fidelity mimicking viral polymerases, as well as the RdRp from SARS-CoV, the virus causing the 2003 and subsequent outbreaks of SARS. Our rationale is that the low fidelity viral-like enzymes would incorporate 2’-F,Me-UTP and stop further replication, while the high fidelity polymerases, typical of host cell polymerases, would not. Experimental proof for termination of the SARS-CoV polymerase catalyzed RNA replication would provide further support for this rationale, indicating that Sofosbuvir or its modified forms will inhibit SARS-CoV-2.

We first carried out DNA polymerase extension reactions with the active form of Sofosbuvir (2’-F,Me-UTP) using Thermo Sequenase as an example of high fidelity, host-like polymerases, and two mutated DNA polymerases which are known to be more promiscuous in their ability to incorporate modified nucleotides, Therminator II and Therminator IX, as examples of viral-like low fidelity enzymes. A DNA template-primer complex, in which the next two available bases were A (Fig. 3), was incubated with either 2’-F,Me-UTP (structure shown in Fig. 2a), or dTTP as a positive control, in the appropriate polymerase buffer. If the 2’-F,Me-UTP is incorporated and inhibits further incorporation, a single-base primer extension product will be produced. By contrast, dTTP incorporation will result in primer extension by 2 bases. After performing the reactions, we determined the molecular weight of the extension products using MALDI-TOF-mass spectrometry (MALDI-TOF MS).

**Fig. 3.**
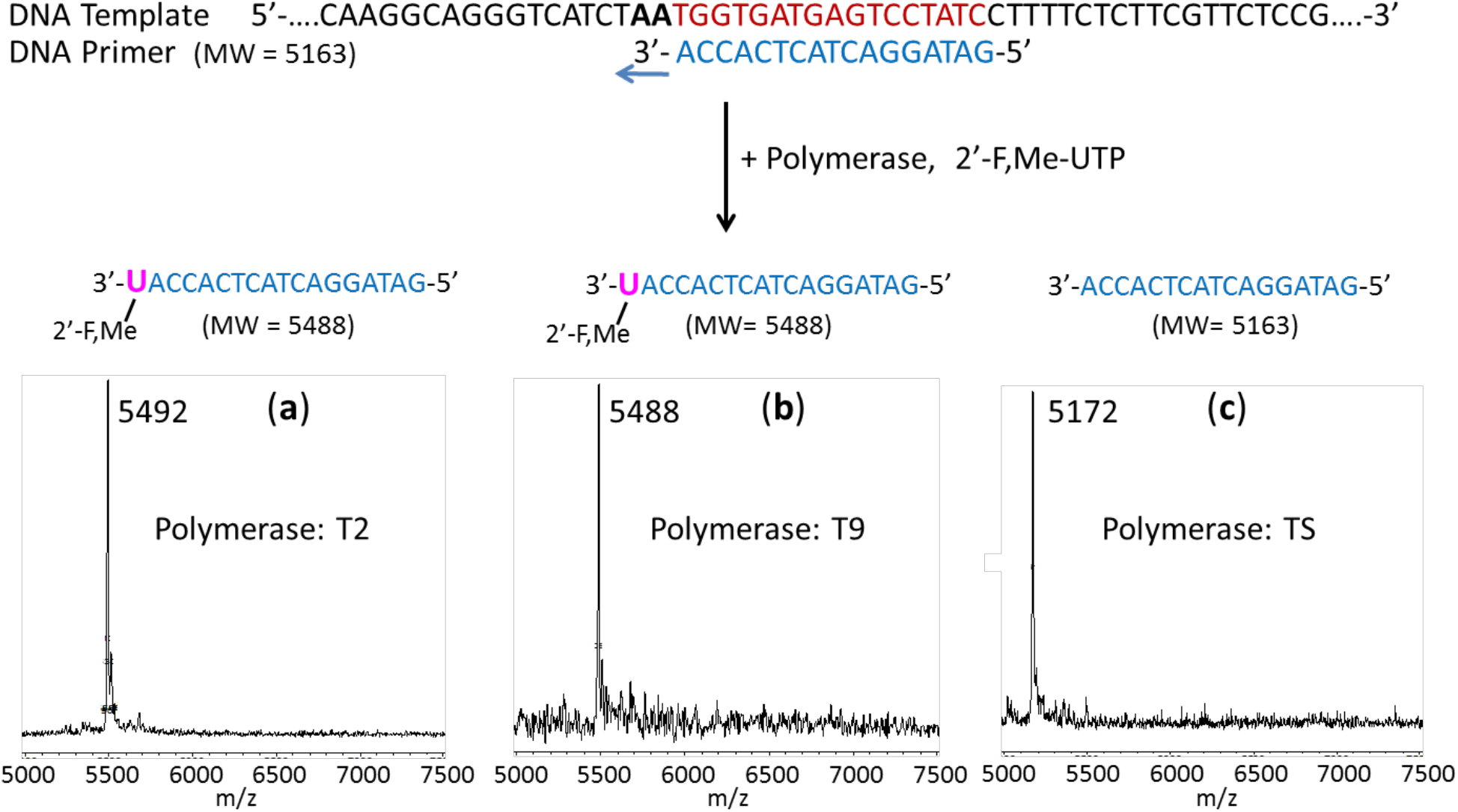
Incorporation of 2’-F,Me-UTP by two low-fidelity polymerases as terminators but not by a high-fidelity polymerase. The sequence of the primer and template used for these extension reactions is shown at the top of the figure. Polymerase extension reactions were performed by incubating the primer and template with 2’-F,Me-UTP and the appropriate reaction buffer for the specific enzyme, followed by detection of the reaction products by MALDI-TOF MS. The MS spectra for Therminator II (T2) in (a) and for Therminator IX (T9) in (b) indicates single-base incorporation and termination, while the MS spectrum for Thermo Sequenase (TS) in (c) indicates no incorporation, showing only a primer peak. The accuracy for m/z determination is ± 10 Da.

As seen in Figs. 3a and 3b, when the primer-template complex (sequences shown at top of Fig. 3) and 2’-F,Me-UTP were incubated with the low fidelity 9°N polymerase mutants,^29–31^ Therminator II (T2) and Therminator IX (T9), we observed single product peaks with molecular weights of 5492 Da and 5488 Da, indicating single base extension in the polymerase reaction. Thus 2’-F,Me-UTP was able to be incorporated and block further nucleotide incorporation. In contrast, when the extension reactions were carried out with high-fidelity Thermo Sequenase DNA polymerase (TS),^32^ there was no incorporation, as evidenced by a single primer peak at 5172 Da (Fig. 3c). This supports our rationale that Thermo Sequenase, a high fidelity enzyme originally designed for accurate Sanger sequencing, will not incorporate 2’-F,Me-UTP, while a low-fidelity polymerase, such as T2 and T9, will incorporate 2’-F,Me-UTP and stop further nucleotide incorporation. When dTTP was used as a positive control with these three enzymes, incorporation continued past the first A in the template, resulting in a higher molecular weight peak.

These results demonstrate that lower fidelity polymerases will have a high likelihood of incorporating 2’-F,Me-UTP and inhibit viral RNA replication, whereas high fidelity enzymes, more typical of the host DNA and RNA polymerases, will have a low likelihood of being inhibited by 2’-F,Me-UTP. Anti-viral drug design based on this principle may lead to potent viral polymerase inhibitors with fewer side effects. To provide further proof that SARS-CoV-2 RdRp might be inhibited by 2’-F,Me-UTP, we next tested the ability of this molecule to be incorporated into an RNA primer to terminate the reaction catalyzed by the RdRp from SARS-CoV, using an RNA template. As shown in Fig. 4a, the active triphosphate form of the drug not only was incorporated by the RdRp, but prevented further incorporation, behaving as a terminator in the polymerase reaction.

**Fig. 4.**
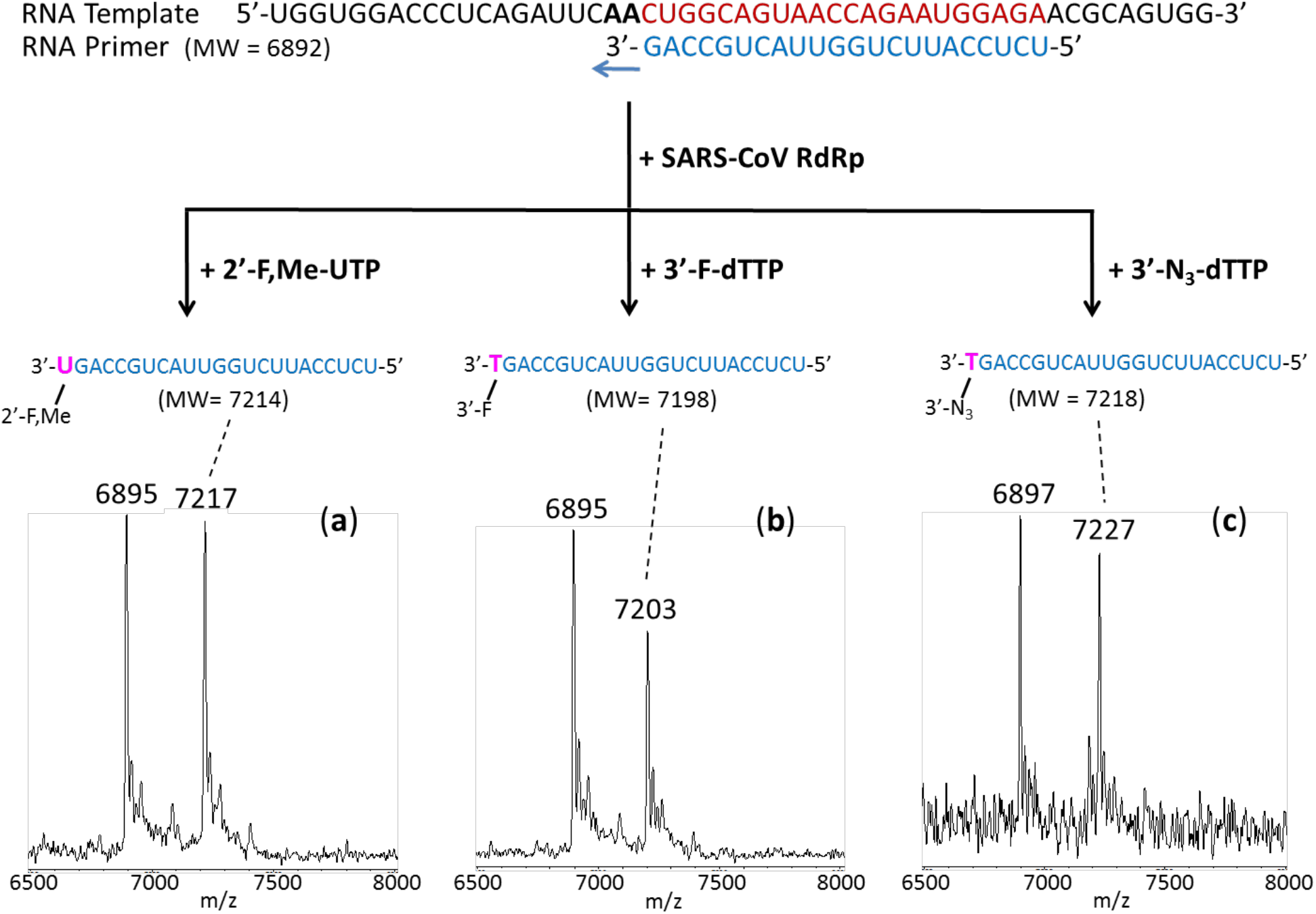
Incorporation of 2’-F,Me-UTP, 3’-F-dTTP and 3’-N_3_-dTTP by SARS-CoV RdRp to terminate the polymerase reaction. The sequence of the primer and template used for these extension reactions, which are within the N1 coding sequence of the SARS-CoV-2 genome, is shown at the top of the figure. Polymerase extension reactions were performed by incubating (a) 2’-F,Me-UTP, (b) 3’-F-dTTP, and (c) 3’-N_3_-dTTP with pre-assembled SARS-CoV polymerase (nsp12, nsp7 and nsp8), the indicated RNA template and primer, and the appropriate reaction buffer, followed by detection of reaction products by MALDI-TOF MS. The detailed procedure is shown in the Methods section. For comparison, data for extension with UTP are presented in Extended Data Fig. 2. The accuracy for m/z determination is ± 10 Da.

Based on our similar insight related to their molecular structures and previous antiviral activity studies, in comparison with Sofosbuvir, we selected the triphosphate forms of Alovudine (3’-deoxy-3’-fluorothymidine) and azidothymidine (AZT, the first FDA approved drug for HIV/AIDS) for evaluation as inhibitors of the SARS-CoV RdRp. These two molecules share a similar backbone structure (base and ribose) to Sofosbuvir, but have fewer modification sites and less steric hindrance. Furthermore, because these modifications on Alovudine and AZT are on the 3’ carbon in place of the OH group, they directly prevent further incorporation of nucleotides leading to permanent termination of RNA synthesis and replication of the virus.

Alovudine is one of the most potent inhibitors of HIV reverse transcriptase and HIV-1 replication. This promising drug was discontinued after a Phase II trial due to its hematological toxicity.^33^ However, subsequent *in vitro* studies showed Alovudine was very effective at suppressing several nucleoside/nucleotide reverse transcriptase inhibitor (NRTI)-resistant HIV-1 mutants.^34^ New clinical studies were then carried out in which low doses of Alovudine were given as supplements to patients showing evidence of infection by NRTI resistant HIV strains and not responding well to their current drug regimen. A 4-week course of 2 mg/day Alovudine reduced viral load significantly, and was relatively well tolerated with no unexpected adverse events.^35^

AZT is another antiretroviral medication which has long been used to prevent and treat HIV/AIDS.^36–38^ Upon entry into the infected cells, similar to Alovudine, cellular enzymes convert AZT into the effective 5’-triphosphate form (3’-N_3_-dTTP, structure shown in Fig. 2d), which competes with dTTP for incorporation into DNA by HIV-reverse transcriptase resulting in termination of HIV’s DNA synthesis.^39^ Since the side effects and toxicity of AZT are well understood, novel methodologies have been directed at enhancing AZT plasma levels and its bioavailability in all human organs in order to improve its therapeutic efficacy. Among these possibilities, an AZT prodrug strategy was proposed.^40^

We thus assessed the ability of 3’-N_3_-dTTP and 3’-F-dTTP, the active triphosphate forms of AZT and Alovudine, along with 2’-F,Me-UTP, to be incorporated by SARS-CoV RdRp into an RNA primer and terminate the polymerase reaction.

The RdRp of SARS-CoV, referred to as nsp12, and its two protein cofactors, nsp7 and nsp8, shown to be required for the processive polymerase activity of nsp12, were cloned and purified as described.^41,42^ These three viral gene products have high homology (e.g., 96% identity and 98% similarity for nsp12, with similar homology levels at the amino acid level for nsp7 and nsp8) to the equivalent gene products from SARS-CoV-2, the causative agent of COVID-19. A detailed description of the homologies of nsp7, nsp8 and nsp12 is included in Extended Data Fig. 1 which highlights key functional motifs in nsp12 described by Kirchdoerfer and Ward.^42^ Of these, Motifs A, B, E, F and G are identical in SARS-CoV and SARS-CoV-2 at the amino acid level, and Motifs C and D display only conservative substitutions.

We performed polymerase extension assays with 2’-F,Me-UTP, 3’-F-dTTP, 3’-N_3_-dTTP or UTP following the addition of an pre-annealed RNA template and primer to a pre-assembled mixture of the RdRp (nsp12) and two cofactor proteins (nsp7 and nsp8). The extended primer products from the reaction were subjected to MALDI-TOF-MS analysis. The RNA template and primer, corresponding to the N1 epitope region of the N protein of the SARS-CoV-2 virus, were used for the polymerase assay, and their sequences are indicated at the top of Fig. 4. Because there are two As in a row in the next available positions of the template for RNA polymerase extension downstream of the priming site, if the 2’-F,Me-UTP, 3’-F-dTTP or 3’-N_3_-dTTP are incorporated by the viral RdRp, a single nucleotide analogue will be added to the 3’-end of the primer strand. If they are indeed inhibitors of the polymerase, the extension should stop after this incorporation; further 3’-extension should be prevented. In the case of the UTP control reaction, two UTP’s should be incorporated. As shown in Fig. 4 and Extended Data Fig. 2, this is exactly what we observed. In the MALDI-TOF MS trace in Fig. 4a, a peak indicative of the molecular weight of a one-base 2’-F,Me-UTP analogue primer extension product was obtained (7217 Da observed, 7214 Da expected). Similarly, in the trace in Fig. 4b, a single extension peak indicative of a single-base extension by 3’-F-dTTP is revealed (7203 Da observed, 7198 Da expected), with no further incorporation. And in the trace in Fig. 4c, a single extension peak indicative of a single-base extension by 3’-N_3_-dTTP is seen (7227 Da observed, 7218 Da expected), with no evidence of further incorporation. As a positive control, extension by 2 UTPs occurred (7506 Da observed, 7504 Da expected) in the MALDI-TOF MS trace in Extended Data Fig. 2.

In summary, these results demonstrate that the nucleotide analogues 2’-F,Me-UTP, 3’-F-dTTP and 3’-N_3_-dTTP, are permanent terminators for the SARS-CoV RdRp. Their prodrug versions (Sofosbuvir, 3’-F-5’-*O*-phosphoramidate dT nucleoside and 3’-N_3_-5’-*O*-phosphoramidate dT nucleoside) can be readily synthesized using the ProTide prodrug approach, as shown in Fig. 2a, c and d, and can be developed as therapeutics for both SARS and COVID-19.

One factor that has confounded the development of RdRp inhibitors in coronaviruses is the presence of a 3’-exonuclease-based proofreading activity such as that associated with nsp14, a key component of the replication-transcription complex in SARS-CoV,^43,44^ and also encoded in SARS-CoV-2. This exonuclease activity can be overcome with the use of 2’-O-methylated nucleotides.^43^ Importantly, since both Sofosbuvir and AZT are FDA approved drugs, where toxicity tests have already been performed, they can be evaluated quickly in laboratory and clinical settings.

We have recently described^2^ a strategy to design and synthesize viral polymerase inhibitors, by combining the ProTide Prodrug approach^17^ used in the development of Sofosbuvir with the use of 3’-blocking groups that we have previously built into nucleotide analogues that function as terminators for DNA sequencing by synthesis.^29,30,45^ We reasoned that (i) the hydrophobic phosphate masking groups will allow entry of the compounds into the virus-infected cells, (ii) the 3’-blocking group on the 3’-OH with either free 2’-OH or modifications at the 2’ position will encourage incorporation of the activated triphosphate analogue by viral polymerases but not host cell polymerases, thus reducing any side effects, and (iii) once incorporated, further extension will be prevented by virtue of the 3’-blocking group, thereby completely inhibiting viral replication and potentially overcoming the development of resistance due to the accumulation of new mutations in the RdRp.^46^

Our design criterion is to identify groups for attachment to the 3’-OH with appropriate structural and chemical properties (e.g., size, shape, rigidity, flexibility, polarity, reactivity [e.g., stability to cellular enzymes]),^47,48^ along with appropriate 2’-substitutions, so that they will be incorporated by the viral RdRp, while minimizing incorporation by the host polymerases. We previously used this chemical and structural principle to select a variety of chemical moieties that block the 3’-OH of the nucleotide analogues as polymerase terminators.^31,49,50^

In conclusion, we demonstrated the capability of more tolerant DNA polymerases, as well as SARS CoV RNA-dependent RNA polymerase, which is nearly identical to the SARS-CoV-2 RdRp responsible for COVID-19, to incorporate 2’-F,Me-UTP, the active form of Sofosbuvir, where it serves to terminate the polymerase reaction. We also showed two other nucleotide triphosphates, 3’-F-dTTP, the active form of Alovudine, and 3’-N_3_-dTTP, the active form of AZT, can also be incorporated and terminate further nucleotide extension by the RdRp in the polymerase reaction, potentially preventing further replication of the virus. Prodrug versions of these compounds and their derivatives therefore can be developed as potent broad-spectrum therapeutics for coronavirus infectious diseases, including SARS, MERS and COVID-19.

## EXTENDED DATA

### Methods

#### Extension reactions with DNA polymerases

Oligonucleotides were purchased from Integrated DNA Technologies (IDT Inc.). The 20 μl extension reactions consisted of 3 μM DNA template and 5 μM DNA primer (sequences shown in Fig. 3), 10 μM 2’-F,Me-UTP (Sierra Bioresearch) or 10 μM dTTP, 1× Thermo Sequenase buffer or 1× ThermoPol buffer (for Therminator enzymes), and either 10 U Thermo Sequenase (GE Healthcare), 4 U Therminator II or 10 U Therminator IX (New England Biolabs). The 1× Thermo Sequenase buffer consists of 26 mM Tris-HCl, pH 9.5 and 6.5 mM MgCl_2_. The 1× ThermoPol buffer contains 20 mM Tris-HCl, pH 8.8, 10 mM (NH_4_)_2_SO_4_, 10 mM KCl, 2 mM MgSO_4_, and 0.1% Triton X-100. Incubations were performed in a thermal cycler using 15 cycles of 30 sec each at 65°C, 45°C and 65°C. Following desalting using an Oligo Clean & Concentrator (Zymo Research), the samples were subjected to MALDI-TOF-MS (Bruker ultrafleXtreme) analysis, following a previously described method.^29^

#### Extension reactions with RNA-dependent RNA polymerase

Oligonucleotides were purchased from IDT, Inc. Following a published strategy,^41,42^ the primer and template (sequences shown in Fig. 4) were annealed by heating to 70°C for 10 min and cooling to room temperature in 1× reaction buffer. The RNA polymerase mixture consisting of 2 *μ*M nsp12 and 6 *μ*M each of cofactors nsp7 and nsp8 was incubated for 15 min at room temperature in a 1:3:3 ratio in 1× reaction buffer. Then 5 *μ*l of the annealed template primer solution containing 2 *μ*M template and 1.7 *μ*M primer in 1× reaction buffer was added to 10 *μ*l of the RNA polymerase mixture and incubated for an additional 10 min at room temperature. Finally 5 *μ*l of a solution containing either 2 mM 2’-F,Me-UTP, 2 mM 3’-F-dTTP or 2 mM UTP in 1× reaction buffer was added, and incubation was carried out for 2 hr at 30°C. The final concentrations of reagents in the 20 *μ*l extension reactions were 1 *μ*M nsp12, 3 *μ*M nsp7, 3 *μ*M nsp8, 425 nM RNA primer, 500 nM RNA template, either 500 *μ*M 2’-F,Me-UTP (Sierra Bioresearch), 500 *μ*M 3’-F-dTTP (Amersham Life Sciences), or 500 *μ*M 3’-N_3_-dTTP (Amersham Life Sciences), and 1× reaction buffer (10 mM Tris-HCl pH 8, 10 mM KCl, 2 mM MgCl_2_ and 1 mM β-mercaptoethanol). [In the experiment with UTP shown in Extended Data Fig. 2, the final concentrations were 500 nM nsp12, 1.5 *μ*M nsp7, 1.5 *μ*M nsp8, 425 nM RNA primer, 250 nM RNA template and 500 *μ*M UTP (Fisher) and the reaction time was 1 h at 30°C.] Following desalting using an Oligo Clean & Concentrator (Zymo Research), the samples were subjected to MALDI-TOF-MS (Bruker ultrafleXtreme) analysis.

## Acknowledgments

This research is supported by Columbia University, which has filed a patent application on the work described in this manuscript.

## Author Contributions

J.J. conceived and directed the project; the approaches and assays were designed and conducted by J.J., X.L., S.K., S.J., J.J.R., M.C. and C.T., comparative sequence analysis was performed by I.M. and S.K., and SARS-CoV polymerase and associated proteins nsp12, 7 and 8 were cloned and purified by R.N.K. Data were analyzed by all authors. All authors wrote and reviewed the manuscript.

---

**Extended Data Fig. 1.**
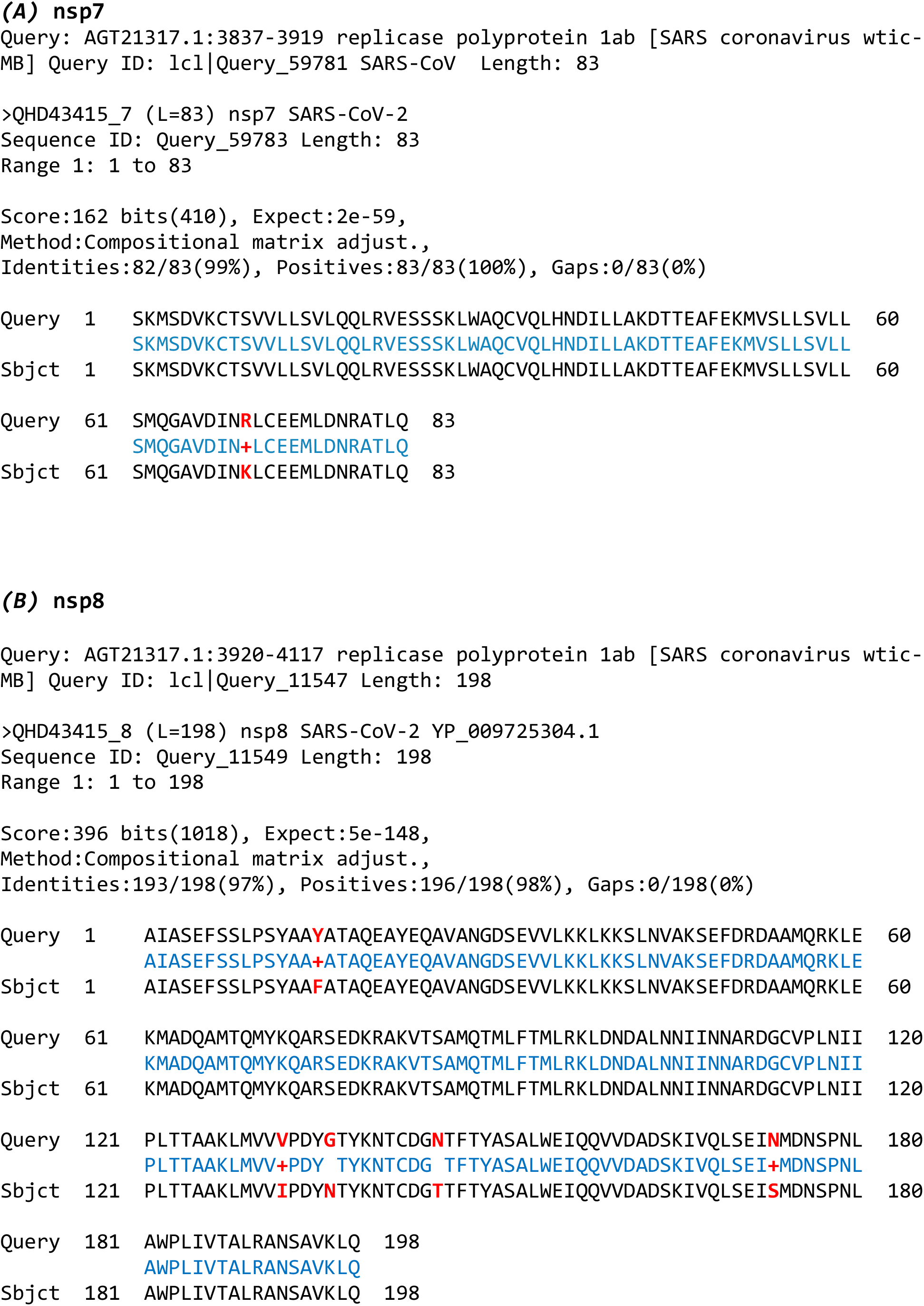
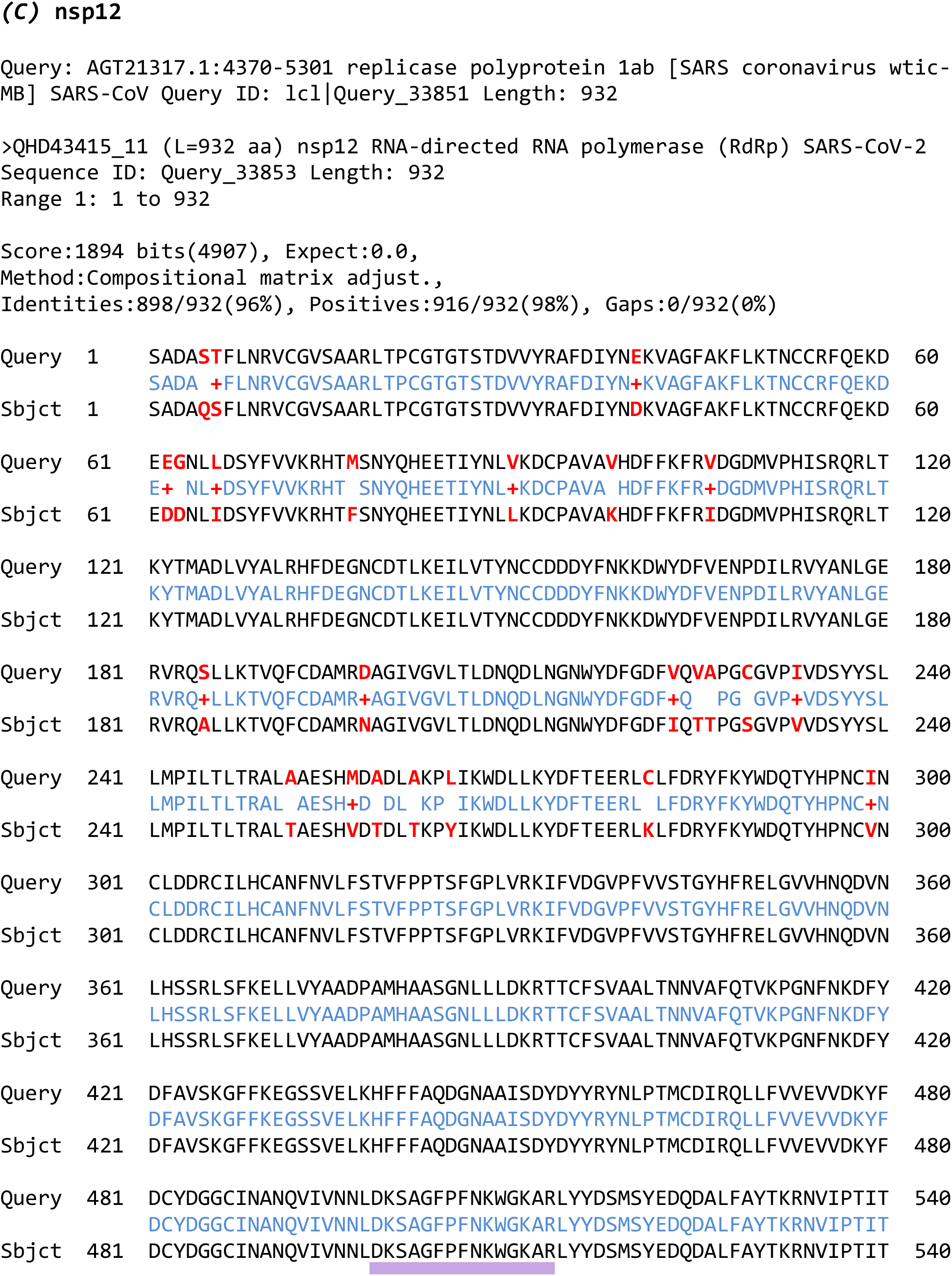
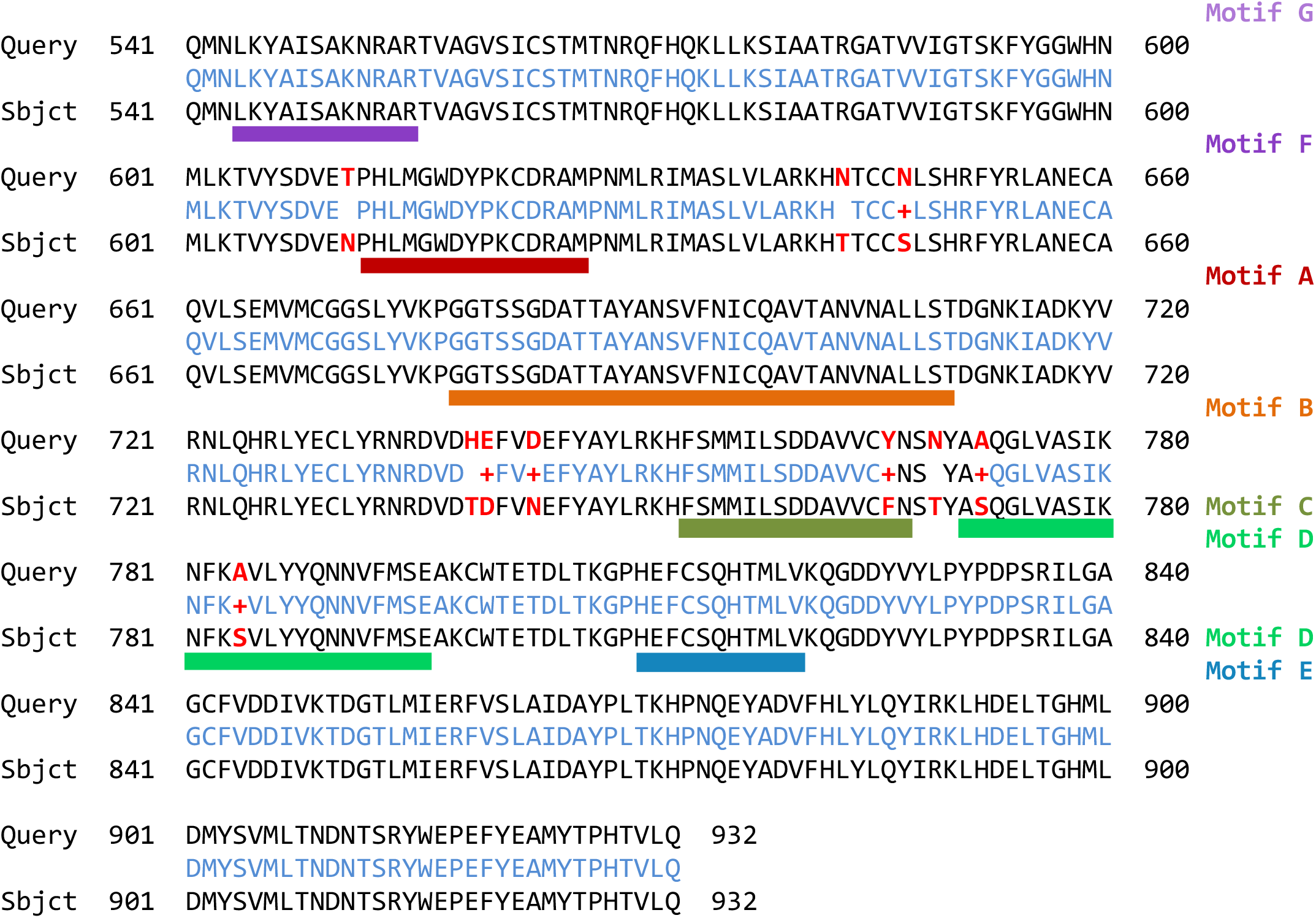
Protein sequence alignments for nsp7, nsp8, and nsp12: SARS-CoV vs SARS-CoV-2. Protein sequences are from NCBI Protein Database, accession ID’s as indicated. Sequences were aligned with blastp.^51^ Consensus is shown in blue between the query and subject sequences; positions of amino acid substitutions are in red; + indicates conservative amino acid substitutions involving amino acids with close physico-chemical properties. ***(A)*** nsp7; ***(B)*** nsp8; ***(C)*** nsp12: functional motifs^42^ are shown as colored bars underneath the aligned sequences. Comparison of the polymerase complex components (nsp7, nsp8, and nsp12 proteins) shows that these proteins are very similar in SARS-CoV and SARS-CoV-2. There are no indels in any of the three protein pairs. There is only one amino acid substitution in nsp7 (99% sequence identity); nsp8 has 5 amino acid changes (97% sequence identity), out of which 3 are between amino acids with similar properties. Alignment of the nsp12 pair shows that 898 out of 932 amino acids (96%) are identical between SARS-CoV and SARS-CoV-2. Eighteen of the substitutions are between amino acids with similar physico-chemical properties, therefore the level of similarity is higher, at 98%. Most of the amino acid substitutions (24 out of 34) are located within the N-terminal portion of the nsp12 protein. This region corresponds to the NiRAN domain (nidovirus RdRp-associated nucleotidyltransferase; approximately amino acids 1 through 250) which is also less conservative in other coronaviruses.^5^ Within the next region (the interface domain, aa ~250 through 400), the first 15 amino acid positions have multiple substitutions, but the rest of the interface domain is quite conservative. The region beyond the interface domain, corresponding to the nsp12 C-terminus, contains polymerase functional domains. These domains constitute the canonical *fingers, palm*, and *thumb* of the polymerase enzyme and contain several motifs that are conservative among coronaviruses (Motifs A through F). Out of the 34 amino acid substitutions in the nsp12 between SARS-CoV and SARS-CoV-2, only three substitutions are located within these motifs, and all three are between similar amino acids.

**Extended Data Fig. 2.**
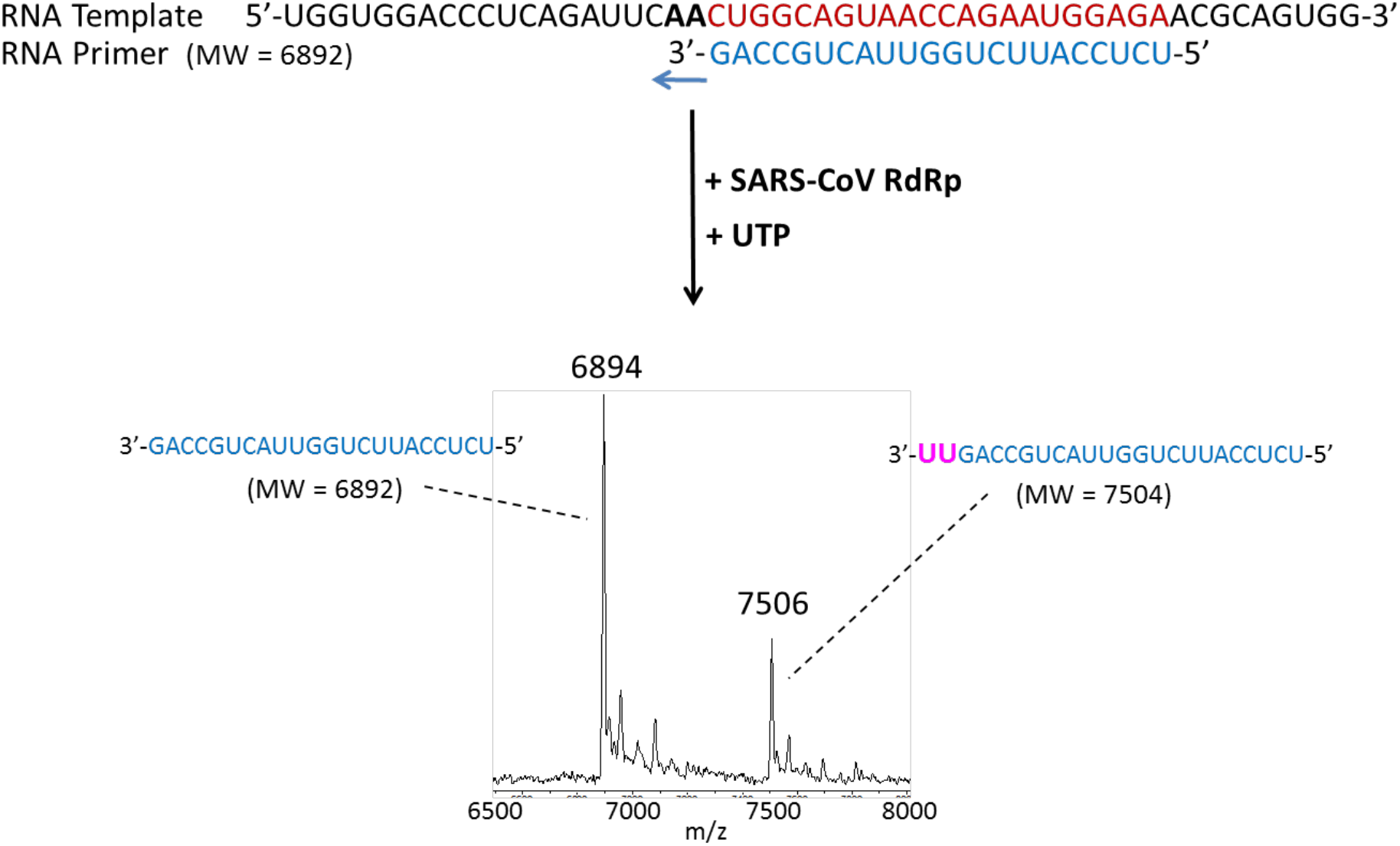
Incorporation of UTP by SARS-CoV RNA-dependent RNA polymerase. The sequence of the primer and template used for this extension reaction is shown at the top of the figure. Polymerase extension reactions were performed by incubating UTP with pre-assembled SARS-CoV polymerase (nsp12, nsp7 and nsp8), the indicated RNA template and primer, and the appropriate reaction buffer, followed by detection of reaction products by MALDI-TOF MS. The detailed procedure is shown in the methods. The accuracy for m/z determination is ± 10 Da.

